# Sequence alignment using machine learning for accurate template-based protein structure prediction

**DOI:** 10.1101/711945

**Authors:** Shuichiro Makigaki, Takashi Ishida

## Abstract

**Motivation:** Template-based modeling, the process of predicting the tertiary structure of a protein by using homologous protein structures, is useful if good templates can be found. Although modern homology detection methods can find remote homologs with high sensitivity, the accuracy of template-based models generated from homology-detection-based alignments is often lower than that from ideal alignments.

**Result:** In this study, we propose a new method that generates pairwise sequence alignments for more accurate template-based modeling. The proposed method trains a machine learning model using the structural alignment of known homologs. It is difficult to directly predict sequence alignments using machine learning. Thus, when calculating sequence alignments, instead of a fixed substitution matrix, this method dynamically predicts a substitution score from the trained model. We evaluate our method by carefully splitting the training and test datasets and comparing the predicted structure’s accuracy with that of state-of-the-art methods. Our method generates more accurate tertiary structure models than those produced from alignments obtained by other methods.

**Availability and Implementation:** https://github.com/shuichiro-makigaki/exmachina

## 1 Introduction

Proteins are key molecules in biology, biochemistry and pharmaceutical sciences. To reveal the functions of proteins, it is essential to understand the relationships between proteins’ structure and function. Proteins that have similar functions are often evolutionarily related; these proteins are called homologs. Revealing homologs and studying proteins’ structure to deduce their function are crucial molecular biology techniques. Protein structures can be determined by experimental means such as X-ray crystallography or nucleic magnetic resonance; protein structures derived in this way are often registered to and accessible in the online Protein Databank (PDB) (wwPDB consortium, 2018). However, despite improvements in experimental methods for determining protein structures, the speed at which amino acid sequences can be revealed has overtaken our ability to ascertain the corresponding proteins’ structures. Therefore, protein structure prediction, that is, the use of computational techniques to generate a tertiary structural model of a given amino acid sequence, remains essential.

Various methods for predicting a protein structure have been proposed and can be briefly classified as either physicochemical (also called *de novo*) simulations or template-free modeling methods. Other methods, called template-based or homology modeling, predict structures based on templates and their sequence alignment to a target protein. Template structures are the structures of homologous proteins, often found with homology detection methods. Currently, template-based modeling methods are the most practical because the predicted models are much more accurate if we can find good templates and protein sequence alignments. In long-term homology detection studies from FASTA (Pearson and Lipman, 1988) and BLAST (Altschul *et al.*, 1990), sequence profiles based on multiple sequence alignments, such as PSI-BLAST (Altschul *et al.*, 1997) and DELTA-BLAST (Boratyn *et al.*, 2012), have detected homology with high accuracy. Hidden Markov model (HMM)-based methods, a subset of sequence profile-based methods, also detect remote homologs; HMM comparison methods, such as HHpred (Zimmermann *et al.*, 2018) have performed excellently in structure prediction benchmarks (Hildebrand *et al.*, 2009; Meier and Söding, 2015).

Recent homology search methods have been able to detect remote homologs, although sometimes sufficiently accurate structure models cannot be obtained because the quality of the sequence alignment generated by homology detection program is poor. If a more accurate model is required, researchers must often edit alignments manually before modeling to improve their quality. In structural alignment, the structural difference between a target protein structure and a template protein structure is minimized; thus, sequence alignments generated by structural alignment are ideal for template-based modeling (Figure 1). Often, the sequence alignments generated by the homology detection methods are dissimilar to those generated by structural alignment, especially for remote homologs. In essence, alignment quality is crucial to template-based modeling. Thus far, a method’s ability to detect remote homologs has been prioritized because models cannot be generated without a template. However, to achieve higher-accuracy template-based modeling, the improvement of sequence alignment generation is a critical open problem.

**Fig. 1.**
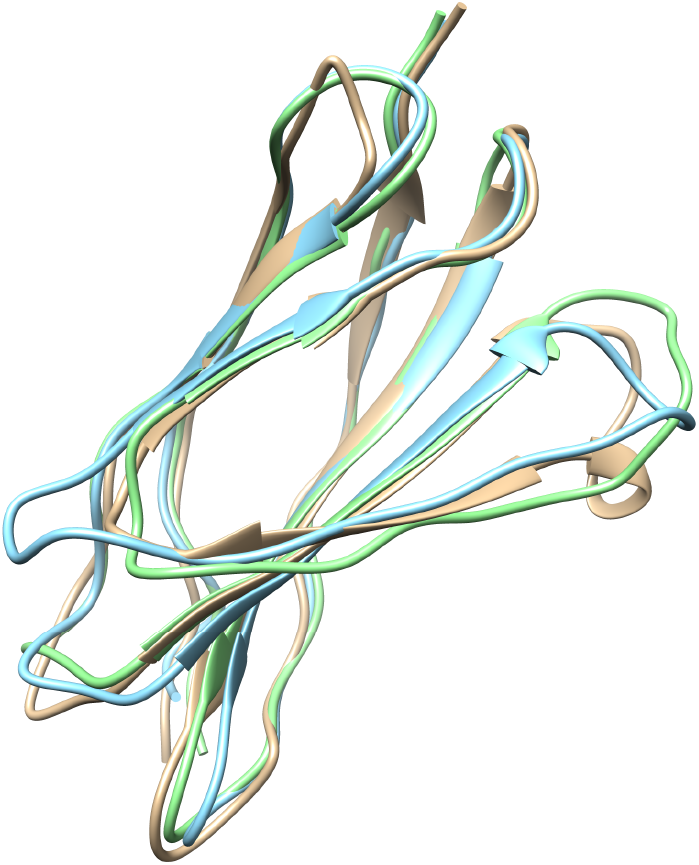
Model differences. Query (yellow) and template proteins are 1QG3A and 1VA9A, respectively. The green model is generated from a structural alignment (TM-align), and the blue model is from HHsearch. The TM-scores of HHsearch and structural alignment are 0.801 and 0.881, respectively [Molecular graphics were performed with the UCSF Chimera (Pettersen et al., 2004) package.]

This problem has been mentioned in several studies (Kopp *et al.*, 2007) in which researchers have tried to improve alignments manually based on their knowledge of biology; fully automated methods are still required. Hijikata *et al.* (2011) proposed an automated method to improve alignments by optimizing gap penalties. They evaluated the premise of gap location in protein 3D structures by examining large protein structure datasets, and found that the distribution of gaps in protein 3D structures differed from previous studies. However, they used the technique mainly for homology detection and the quality of its alignments for prediction model accuracy was still unclear.

Recently, machine learning methods have demonstrated power in homology detection, fold recognition, residue contact map prediction, dihedral prediction, model quality assessment and secondary structure prediction (Cao *et al.*, 2016; Lyons *et al.*, 2014; Manavalan and Lee, 2017; Wang *et al.*, 2017, 2016; Wei and Zou, 2016). Machine learning also seems effective for tackling the problem of alignment generation for homology modeling. However, this topic has not been studied because it is difficult to treat alignment generation as a classification or regression problem.

In this paper, we propose a new pairwise sequence alignment generation method based on a machine learning model that learns the structural alignments of known homologs. Because it is difficult to directly predict sequence alignment using machine learning, we instead use dynamic programming during sequence alignment to dynamically predict a substitution score from the learned model instead of a fixed substitution matrix or profile comparison. Machine learning is used in this substitution score prediction process. We evaluate the proposed method using a carefully split training and test dataset and compare the accuracy of predicted structure models with those of state-of-the-art methods as a measure of sequence alignment quality.

## 2 Materials and methods

Generally, sequence alignment generation is integrated with the homology detection process and the detection tools output sequence alignments with homology search results from the database. In this study, we focus only on alignment generation. Thus, the inputs are a target’s amino acid sequence (query) and another amino acid sequence that was detected as a template by any homology detection method (subject), and the output is an alignment that is more suitable for homology modeling. This process is often called re-alignment. Figure 2 shows an overview of our method. The proposed method accepts query and subject amino acid sequences as input, then aligns their sequences using the Smith–Waterman algorithm (Smith and Waterman, 1981). In classical dynamic programming, a substitution matrix such as BLOSUM62 or PAM250, is used to evaluate the match between residue pairs. To improve alignment accuracy, profile comparison methods, including FORTE (Tomii and Akiyama, 2004) and FFAS (Rychlewski *et al.*, 2008), use the similarity between two position-specific score matrices (PSSMs) of a target residue pair. In contrast, we evaluate residue matches based on a supervised machine learning technique. We train a prediction model using pairwise structural alignments of structurally similar protein pairs as the labels of a training dataset. Thus, the method is expected to output similar sequence alignments by structural alignment. The PSSMs of two input sequences are used as input to the prediction model; to predict the match of a residue pair, PSSMs around target residues within a fixed size window are used. Finally, the method returns a sequence alignment and an alignment score as output.

**Fig. 2.**
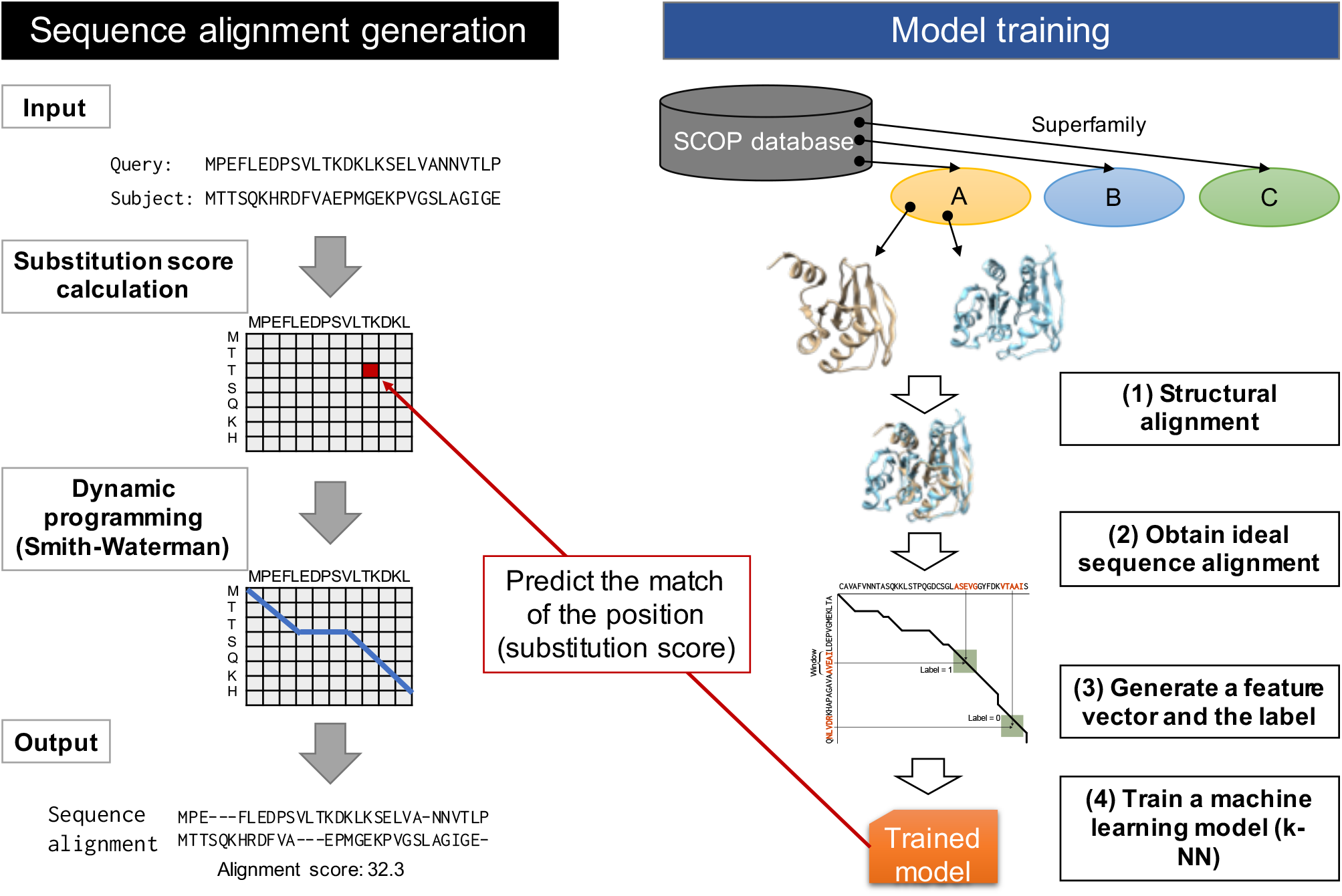
Overview of the proposed method. Two sequences are aligned using the Smith–Waterman algorithm and substitution scores used in the process are estimated by a prediction model. The prediction model is trained to output an alignment similar to the structural alignment

### 2.1 Datasets

Our method needs information about known structurally similar proteins to create structural alignments, for which we used the Structural Classification of Proteins (SCOP) (Fox *et al.*, 2013; Murzin *et al.*, 1995) database. The SCOP database classifies proteins by class, folds, superfamily (SF), family and domain based on manually curated function/structure classifications and contains redundant sequences. Thus, we used the SCOP40 database instead, which contains only domains whose sequence identity is <40% to avoid overfitting and reduce execution time. In this study, we define domains that are in the same SF as structurally similar.

For accurate evaluation and parameter optimization, we split training, test and validation datasets from the full dataset. We selected five domains each from seven SCOP classes to cover various protein structure types, selecting test domains only from SFs containing greater than ten domains. We ignored any small SFs and sorted the remaining domains by their PDB revision date, ultimately selecting 35 domains as test data. For our validation dataset, we split two groups from the remaining dataset. For one group, we selected one domain each from seven SCOP classes for parameter search; the other contains one domain each from the classes for a gap penalty search. Finally, we split 49 domains (= 35 + 7 + 7) from all the datasets for test and validation, and the remaining domains were used for training (see Supplementary Table S1–S3 for details).

In the training dataset, we generated structural alignments of every domain pair in the same SF using TM-align (Zhang and Skolnick, 2005). We treated domain pairs whose TM-align score [TM-score (Zhang and Skolnick, 2004)] was <0.5 as having low structural similarity and filtered them out (Xu and Zhang, 2010). If the SF had only one domain, it was ignored because we could not define a pairwise alignment for it. Finally, 140889 pairwise structural alignments were generated. For PSSM generation, we used three-iteration PSI-BLAST with the UniRef90 (The UniProt Consortium, 2016) database. When the training dataset became too large to process within a reasonable computation time, the training dataset was reduced to 1/10 of its initial size by random selection.

### 2.2 Input vector and label definition

To use machine learning methods to predict matching scores, we had to encode information about residue pairs in a numerical vector representation. In addition, we dealt with the problem as a binary classification problem and used the reliability score of a prediction as a matching score, because structural alignment can only tell us whether a position in a dynamic programming matrix is a match; defining a correct matching score is difficult. Figure 3 shows an overview of this design.

**Fig. 3.**
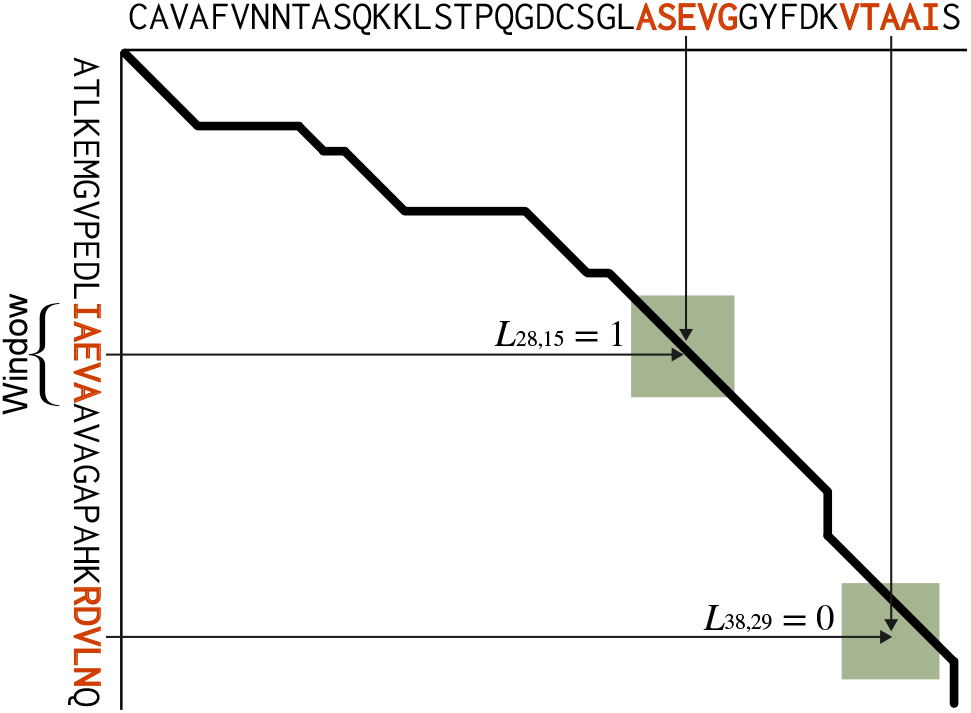
Overview of a feature vector encoding scheme. The *X* and *Y* axes show an amino acid sequence. The bold black line shows the structural alignment path between the sequences on the *X* and *Y* axes, with the green rectangle indicating the window. The feature vector set is calculated only within this window. The feature vector is the concatenation of the PASSM columns of the window subsequence. If the current column is on the line, the label is 1; otherwise, it is 0

Let (*Q, T*) be the query and target sequences, respectively. Let *Q_i_* be the *i*th residue of sequence *Q* and *T_i_* be the *i*th residue of sequence *T*. To encode amino acid sequences in a numerical vector, we make PSSMs of the sequences in advance. The column length of a PSSM is 20, which is the number of amino acid types, and the row length is the length of the sequence. Feature vector **V**_*x,y*_ at *Q_x_* and *T_y_* is the concatenation of the query and target residues’ feature vectors:

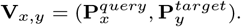

**P**is the concatenation of PSSM rows around the residue, defined as

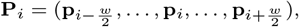

where *w* is the window size and **p**_*i*_ is the *i*th row of the PSSM. Regarding ‘padding’ regions defined in *i* ≤ 0, |*Q*| > *i* and |*T*| > *i*, we assign **p**_*i*_ to be **0**. For example, in the case of *w* = 5, the feature vector dimension is 200 = 20 × 5 × 2.

We can define this feature vector at every residue pair of the query and target sequences. However, we calculate them only within areas where the window moves along with the alignment path because information from residue pairs that are far from the alignment path is not informative.

We assign label *L_x,y_* at *Q_x_* and *T_y_* to be 0 or 1:

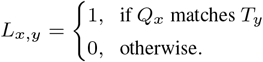

The inputs are a pair of query and template PSSMs and a residue position. The outputs are a predicted label and the normalized confidence score (0 ≤ score ≤ 1).

### 2.3 Alignment calculation

The pairwise sequence alignment of input sequences is calculated using the Smith–Waterman algorithm (Smith and Waterman, 1981), which requires a substitution score for each residue pair. We predict this score using supervised machine learning and the feature vector defined above. Specifically, we used the *k*-nearest neighbor (*k*NN) classification model because it is simple and powerful, especially for large training datasets (Wu *et al.*, 2008). *k*NN calculates the distance between an input feature vector **V**_*x,y*_ and feature vectors in training dataset 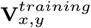 and checks the labels of the *k* nearest feature vectors. Generally, the most major label is output as the predicted label from *k*NN. In this case, 0 or 1 is output because this is a binary classification problem. However, predicting binary labels is too coarse-grained for alignment generation. Thus, the classification confidence score of the *k*NN algorithm, which is the ratio of a predicted positive label, was used as the substitution score of *Q_x_* and *T_y_*, instead.

### 2.4 Parameter optimization

Our method requires some hyperparameters, which we optimized using the validation dataset. We set the number of nearest neighbors to (10, 100, 1000), the gap open penalty to (−0.0001, −0.001, −0.01, −0.1, −1), and the gap extend penalty to (−0.00001, −0.0001, −0.001, −0.01, −0.1, −1). Using a grid search, we selected 1000 as the number of *k*NN neighbors. The affine gap penalty optimizations were −0.1 for gap-open and −0.0001 for gap-extend (Figure 4). These gap penalties were much smaller than the general gap penalties used in other studies because our method’s predicted substitution scores were too small for general gap penalties.

**Fig. 4.**
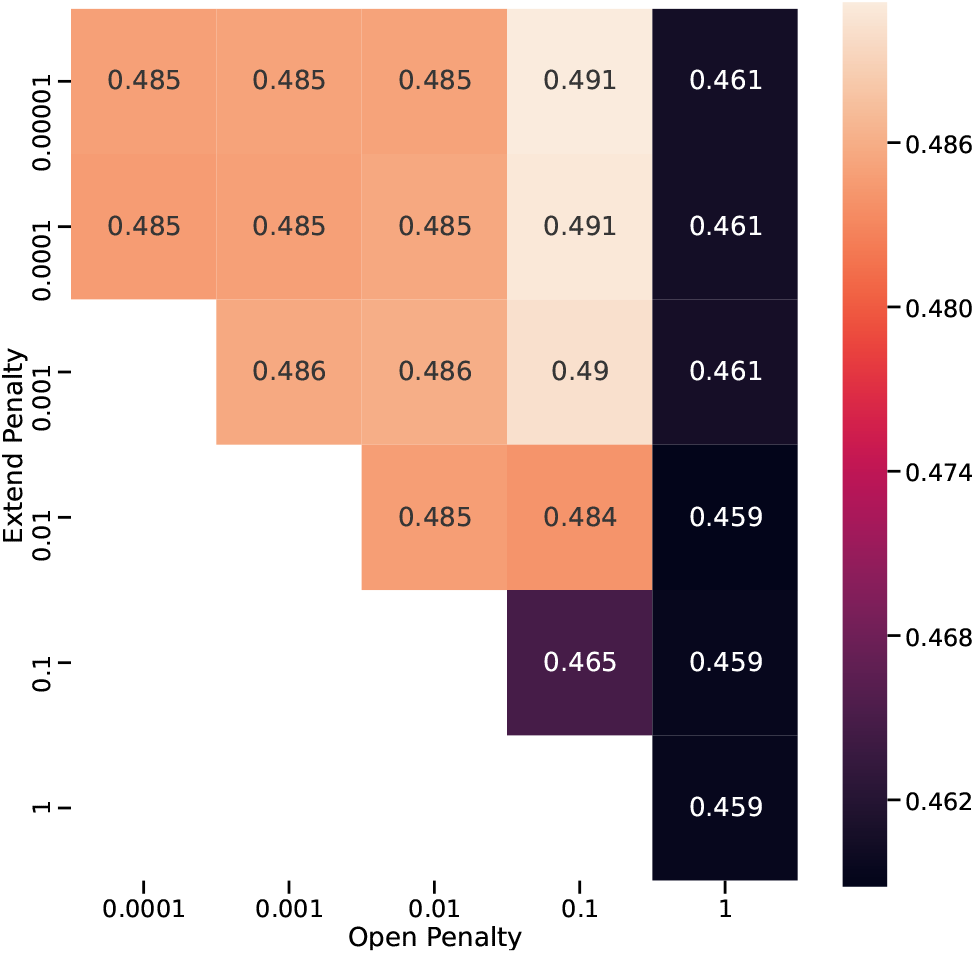
Result of gap penalty grid search. Values in the heatmap show the average TM-score of the validation dataset. Penalties were optimized to −0.1 for gap-open and −0.0001 for gap-extend

## 3 Results

In our method, residue matches at two sequence positions is estimated by *k*NN. It can be considered as an independent binary classification problem. Thus, we first checked the performance of the label prediction process using the receiver operating curve (ROC) and the area under the ROC (AUC). Figure 5 shows the results. The proposed method predicted labels accurately, except for d2axto1, which showed an almost random prediction.

**Fig. 5.**
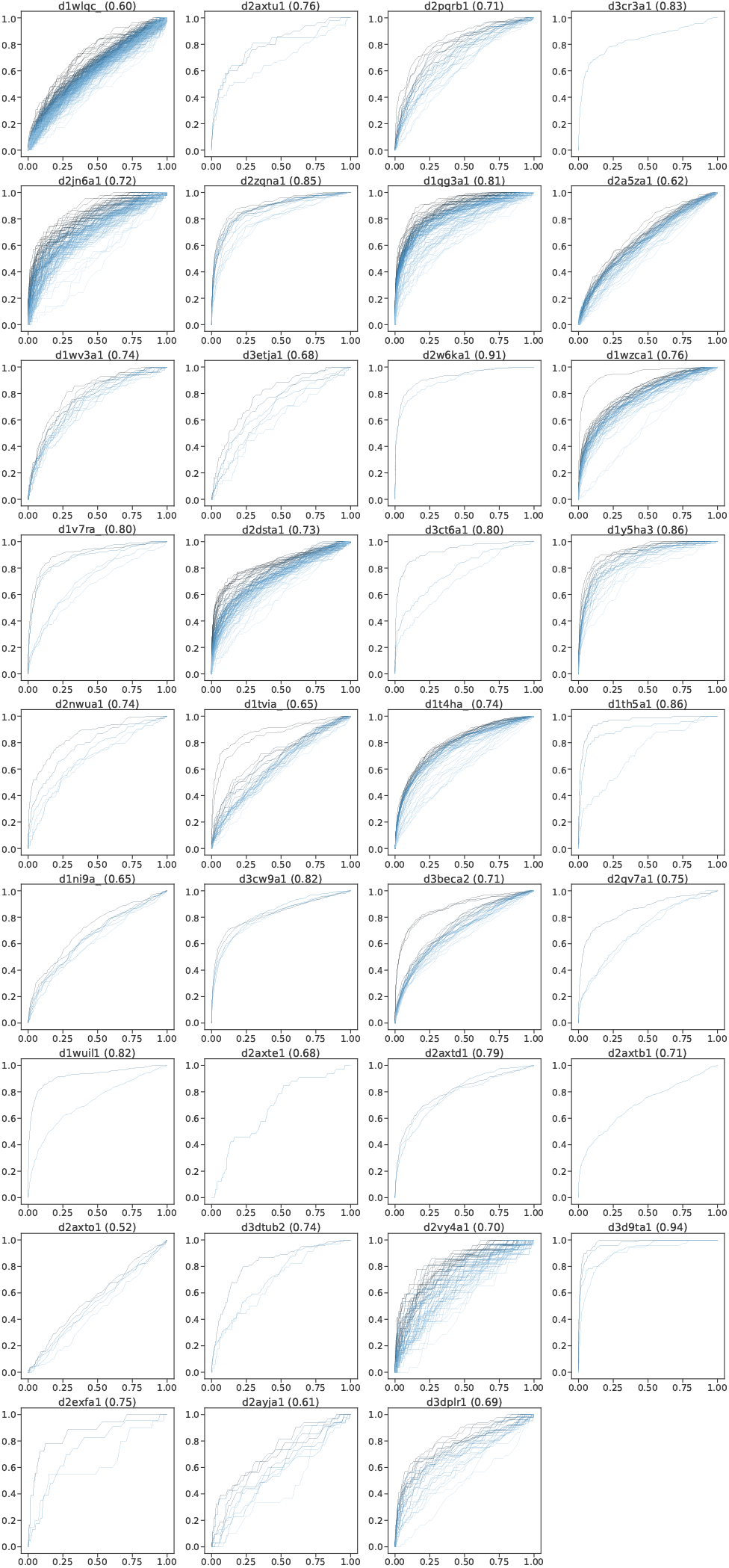
ROC of label prediction. The title is the target name shown in Supplementary Table S1; the average AUC is shown next to the name

Next, we compared the accuracy of three-dimensional predicted protein models generated from these alignments to evaluate the quality of generated sequence alignments. This step is required because there may not be strong correlation between match prediction and model accuracy and we cannot compare our method with other methods directly. We used MODELLER as a modeling tool (Šali and Blundell, 1993). We treated the entire SCOP40 domains, in which sequence similarities in an SF are <40%, in the SF where the query is as a structurally similar protein and applied the proposed method to them. Model accuracy can be evaluated by calculating the similarity between an experimentally resolved structure and a predicted structure. For this purpose, we used TM-score (Zhang and Skolnick, 2004), which evaluates model accuracy by scoring from 0.0 (least accurate) to 1.0 (most accurate).

We used all domains in the SF of the query as template proteins and generated pairwise alignments. We then compared the accuracy of the proposed method with those of PSI-BLAST, DELTA-BLAST, HHsearch (Söding, 2005), the Smith–Waterman algorithm with a BLOSUM62 substitution matrix, and structural alignment. For PSI-BLAST, which accepts a profile as a query, we made profiles by running three iterative PSI-BLAST searches in the UniRef90 database. DELTA-BLAST allows us to use a sequence as a query because it finds profiles from the Conserved Domain Database (Marchler-Bauer *et al.*, 2010) before searching. For HHsearch, we used Uniclust20 (Mirdita *et al.*, 2017) to generate query profiles. We used TM-align (Zhang and Skolnick, 2005) for structural alignment.

Figure 6 shows the accuracy of protein structure prediction. As expected, structural alignments generated the most accurate models (0.551 on average), although the proposed method achieved results that were nearly as accurate (0.499). The naïve Smith–Waterman algorithm and HHsearch performed the next best; their average scores were 0.432 and 0.472, respectively. From the results of data density, in all methods—including proposed method—these results had two peaks. The top-ranking models of all methods showed similar accuracy, but the worst models’ accuracies improved when using the proposed method.

**Fig. 6.**
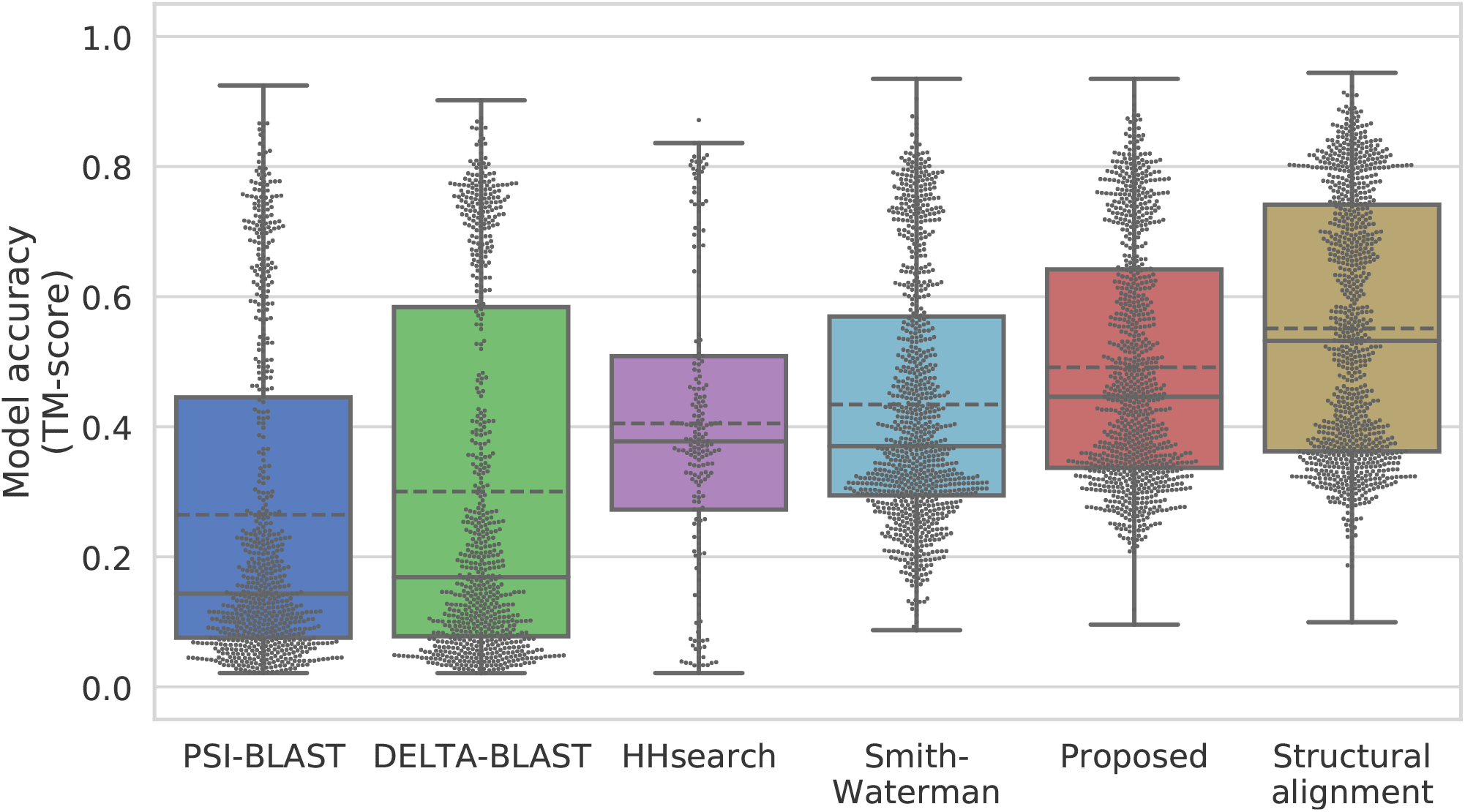
Sequence-independent TM-score of the proposed and competitor methods. The solid line shows the medians and the dashed line shows the means. Dots indicate the data density at each TM-score

In figure 7, we show as an example one of the generated models and an actual alignment, indicating that the proposed method could improve model accuracy. The proposed method succeeded in aligning almost a whole protein and generated very similar alignment results to the structural alignment method. In contrast, HHsearch failed to correctly align a region around the 4th beta strand (residue numbers 40–55) and caused structural differences from the native structure in the loop regions on both sides of the sheet. Figure 7 shows how the proposed method correctly aligned the region that HHsearch failed to align. The *k*NN predicted match score between a query residue with residue number 49 and a template residue with residue number 63 is much higher than the scores around the position. Thus, the proposed method generated an alignment passing through the position.

**Fig. 7.**
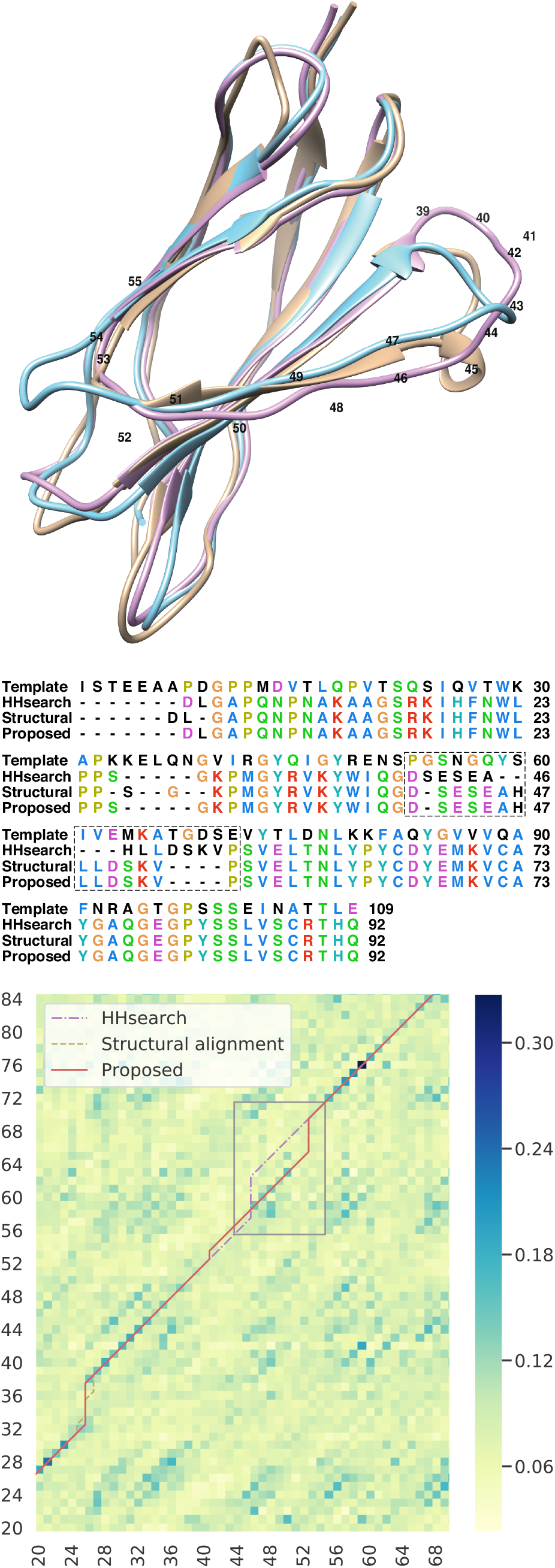
The yellow model in the top figure represents the native structure, the red model is generated by the proposed method, and the blue model is from HHsearch. The TM-scores of HHsearch and our method are 0.815 and 0.871, respectively. The bottom figure is an excerpt of score heatmap and alignment paths. *X* and *Y* axes show the query (1QG3A) and template (1VA9A) residue numbers, respectively. HHsearch (dotted dash) generated different alignments between #46 and #55 from the structural alignment (dash), whereas the proposed method (solid) could generate similar alignments to the structural alignment

## 4 Discussion

### 4.1 Impact of model accuracy improvement for protein function estimation

As shown in our results, we achieved improved model accuracy in structural similarity to a native structure. However, it is difficult to judge whether this improvement is useful for advanced applications, such as protein function estimation. The protein shown in Figure 7 is fibronectin type Ⅲ domain of integrin *β*4, which makes a complex with plectin’s actin-binding domain (Song *et al.*, 2015). Thus, we applied a protein–protein docking to the modeled structures and a ligand structure from the complex structure (PDB ID: 4Q58, Chain: A), using MEGADOCK 4.0 (Ohue *et al.*, 2014) with the default settings, to check the influence of model accuracy. Figure 8 shows the docking results using the model from the proposed method and that from HHsearch.

**Fig. 8.**
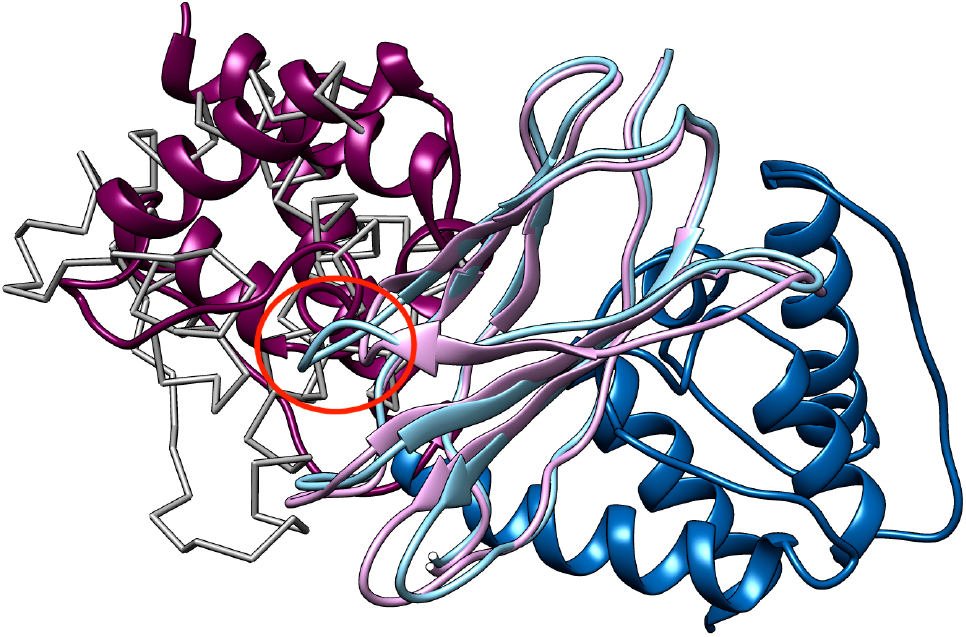
Docking results using modeled structures. The light purple model is generated by the proposed method and the light blue model is from HHsearch. The dark purple model is a ligand structure (plectin) docked by MEGADOCK using the model from the proposed model (ninth model, LRMSD=8.8Å), and the dark blue model is a ligand structure docked using the model from HHsearch (fourth model, LRMSD=23.8Å). The light gray model drawn by C_*α*_ trace shows the correct position based on the native complex structure. The red circle shows a loop that HHsearch failed to model correctly

The best docking model selected from the top-10 models is shown. Where the docking calculation based on the HHsearch model failed to detect the correct binding position, the docking calculation based on the proposed method’s model succeeded. In the HHsearch model, a loop region after 4th *β* strand (the red circle in Figure 8) became longer than the correct structure because of this wrong alignment, causing a steric clash with the ligand structure. As a result, the docking calculation failed to detect the correct binding position and there was no docked result with ligand root mean squared deviation (LRMSD) < 10Å within 3000 models output by MEGADOCK. This is simply one example, but it indicates that the model accuracy improvement achieved with the proposed method is sometimes effective in aid our understanding of the function of a protein.

### 4.2 Application for homology detection

Our method can be used for homology detection by sorting the alignment scores it includes in its result. We investigated the method’s homology detection and the top model accuracies of a search result ranking. The proposed method’s homology detection performance was compared with those of PSI/DELTA-BLAST and HHsearch, as shown in Table 1. To ensure a reasonable computation time, the training dataset was reduced to 1/100 instead of 1/10. ROC_*n*_ considered results only up to the *n*th false positive and AUC_*n*_ was regularized by the number of false positives and cutoff *n*. In this evaluation, we defined true positives as those having the same detected SF as the query and false positives as those having different SFs. Compared with PSI/DELTA-BLAST and HHsearch, the detection sensitivity of the proposed method was lower. The highest average AUC_50_ of HHsearch was 0.706. By contrast, the proposed method had the lowest score, 0.205. We think this is because the proposed method shows many false positive results.

**Table 1.** Average AUC_50_ and model accuracy (TM-score) of the proposed and competitor methods

|  | PSI-BLAST | DELTA-BLAST | HHsearch | Proposed |
| --- | --- | --- | --- | --- |
| AUC<sub>50</sub> | 0.323 | 0.340 | 0.706 | 0.205 |
| TM-score | 0.278 | 0.324 | 0.205 | 0.298 |

Using the search results, we applied template-based modeling to the top 10 search result and made 3D models; the models’ accuracy is mentioned in the second row of table 1. The proposed method achieved the second-highest average TM-score, 0.298. From these results, it is difficult to use our method for homology search. Therefore, we consider that our proposed method is currently useful for the alignment generation phase of template-based modeling, after template detection.

### 4.3 Optimization of window size and the influence of training data reduction

We tested our method using (1, 3, 5) as window size candidates and compared the label prediction accuracy using AUC. The results of window sizes (1, 3, 5) were 0.640, 0.689 and 0.701, respectively. We also tested data reduction ratios of (0.001, 0.01, 0.1) and compared the label prediction accuracy using AUC. The results for ratios of (0.001, 0.01, 0.1) were 0.635, 0.676 and 0.701, respectively. Although increasing the window size and reduction ratio may increase accuracy, we could not evaluate them because the size of the required training dataset would be bigger than our computing resources can manage.

## 5 Conclusion

In this paper, we proposed a new sequence alignment generation method that uses machine learning to accurately predict protein structures. Instead of a fixed substitution matrix, the proposed method predicts substitution scores at each residue pair. To apply machine learning, we developed a method that converts pairwise alignments to numerical vectors of latent space, which enables us to employ a supervised machine learning algorithm for sequence prediction. The predicted scores are directly used to generate alignments, which are in turn used as input for template-based modeling. We evaluated the model accuracy of our alignment generation method and found that it outperformed the state-of-the-art methods. We also investigated our method’s ability to detect remote homologies; using AUC_50_ for comparison, our method did not perform better than other methods. However, we found that the proposed method generated relatively accurate 3D models compared with other methods.

Currently, our method requires a long execution time because of the *k*NN algorithm and dataset size. These factors caused us to reduce the amount of training data used because the model’s execution time depends on the number of target proteins as well as protein size. It would be a natural extension of this work to employ faster *k*NN algorithms, including approximate schemes, because our method does not require precise solutions. The proposed feature vector design can be treated as two-dimensional; in the future, we will also consider the use of higher-performance models such as convolutional neural networks.

## Author contributions

SM conducted the computational experiments and wrote the manuscript. TI supervised the study and wrote the manuscript. Both authors have read and approved the final manuscript.

