## Supplementary Tables for "Sequence alignment using machine learning for accurate template-based protein structure prediction"

Table 1 We selected 14 domains from SCOP40 as test data. Domain IDs shown are the SCOP sid numbers.

| Class | Domains |
|---|---|
| a: All alpha proteins | d1wlqc_, d2axtu1, d2pqrb1, d3cr3a1, d2jn6a1 |
| b: All beta proteins | d2zqna1, d1qg3a1, d2a5za1, d1wv3a1, d3etja1 |
| c: Alpha and beta proteins (a/b) | d2w6ka1, d1wzca1, d1v7ra_, d2dsta1, d3ct6a1 |
| d: Alpha and beta proteins (a+b) | d1y5ha3, d2nwua1, d1tvia_, d1t4ha_, d1th5a1 |
| e: Multidomain proteins | d1ni9a_, d3cw9a1, d3beca2, d2qv7a1, d1wuil1 |
| f: Membrane and cell surface proteins | d2axte1, d2axtd1, d2axtb1, d2axto1, d3dtub2 |
| g: Small proteins | d2vy4a1, d3d9ta1, d2exfa1, d2ayja1, d3dplr1 |

Table 2 We selected 7 domains from SCOP40 as validation dataset 1. These are used for hyper-parameter optimization.

| Class | Domains |
|---|---|
| a: All alpha proteins | d2ij2a1 |
| b: All beta proteins | d3d85d1 |
| c: Alpha and beta proteins (a/b) | d3etja2 |
| d: Alpha and beta proteins (a+b) | d2iiza1 |
| e: Multidomain proteins | d1wuis1 |
| f: Membrane and cell surface proteins | d3dhwa1 |
| g: Small proteins | d3d4ub1 |

Table 3 We selected 7 domains from SCOP40 as validation dataset 2. These are used for gap penalty optimization.

| Class | Domains |
|---|---|
| a: All alpha proteins | d1tw9a1 |
| b: All beta proteins | d3e5ua2 |
| c: Alpha and beta proteins (a/b) | d1xria_ |
| d: Alpha and beta proteins (a+b) | d2jmua1 |
| e: Multidomain proteins | d2zd1b1 |
| f: Membrane and cell surface proteins | d2zfga1 |
| g: Small proteins | d2vuti1 |
